# Using GPT-4 to Automate the Generation of Lay Summaries for Cancer Publications

**DOI:** 10.64898/2025.12.11.693290

**Authors:** Emma Purdie, Tony Yu, Jochen Weile, Diana Lemaire, Melanie Courtot

**Affiliations:** Ontario Institute for Cancer Research, Toronto, Ontario, Canada; Medical Biophysics Department, University of Toronto, Toronto, Ontario, Canada; Department of Computer Science, University of Toronto, Toronto, Ontario, Canada

## Abstract

**Background:** Cancer research literature is often riddled with technical jargon that is not digestible to the average person. Individuals interested in research studies may want to contribute through patient partner engagement or sample donation but find the relevant literature overwhelming. Through the generation of lay summaries, previously inaccessible research papers become easier to comprehend, especially for patient partners or data donors. With large language models (LLMs) continuing to advance, so does their capability to summarize large texts.

**Objectives:** In this study, we examined whether LLMs can produce lay summaries of scientific literature at-scale, while maintaining readability and accuracy to their source texts.

**Methods:** We developed a tool to generate lay summaries of open-access article abstracts and their full texts with GPT-4-Turbo. Prompt development aimed for a target 8th grade reading level assessed with Flesch-Kincaid Grade Level. Human-review metrics were used to evaluate readability and accuracy when generated using abstracts versus full text articles.

**Results:** The average Flesch-Kincaid Grade Level Score was 7.13 for abstract-based summaries and 7.39 for full text-based summaries, indicating summaries at around 7th grade reading level. Human-review metrics showed these summaries were of similar readability and accuracy when generated using abstracts versus full text articles, with mean accuracy scores from human review of 7.09 vs 7.42 out of 10 respectively. Additionally, qualitative patient-based assessment indicated these summaries would encourage participation in research studies.

**Conclusion:** By generating lay summaries for complex and lengthy research papers, their scientific information becomes accessible to a larger audience, including patient partners interested in contributing to cancer research. Summaries that are easy to understand will allow participants to make informed decisions about their involvement and appreciate the impact of their contributions if and when their results are published.

**Lay Summary:** This study explores if artificial intelligence (AI) can help make hard to read cancer research papers easier to understand for members of the public.

*Problem:* When people donate cancer tissue samples or participate in research studies, they often want to know how their contributions are being used. However, scientific papers are full of technical language that’s hard for most people to grasp. People in past studies have said this can make them less willing to take part in research.

*Methods:* The study created a computer program using AI (GPT-4-Turbo) to turn complex kidney cancer research papers into simple summaries. They tested whether the AI could summarize both short abstracts and full-length papers effectively. They aimed for summaries at a 6th to 8th-grade reading level. This was to follow Canadian and U.S. health communication guidelines.

*Results:* The AI created 106 summaries. Computer measures showed the summaries were close to a 7th-grade reading level. Though, researchers had to tell the AI to write for a 2nd-grade audience to achieve this. Of note, summaries from short abstracts were just as accurate and readable as those from full papers. Eighteen volunteers, including five patient partners, reviewed the summaries and rated them for clarity and accuracy. They were rated at around 7 out of 10 points. All patient partners said these summaries would help them decide whether to join research studies and feel more informed about how their contributions matter.

*Why It Matters:* This tool could help patients and donors better understand research without needing a science degree. When people can see how studies work, they are more likely to participate in future research. While patient partners emphasized the need for summaries to be accurate and reliable, this approach shows promise as a unique strategy to better connect the public with research.

## Introduction

In order to drive advancements in research, diagnosis, and patient outcomes, recruiting large cohorts of human participants into studies and trials is critical ^1^. However, in our experience with large-scale initiatives such as the International Cancer Genome Consortium (ICGC), participants have reported finding it hard to understand the complex information related to studies and the impact their contribution makes, such as how their donated samples and data derived from them are being used. This gap in understanding of information has been associated with a perception of distrust in research, an identified barrier to participation which decreases patients’ willingness to engage further or enroll in new studies^2^. The current state-of-the-art for research result dissemination is through scientific publications in established academic journals. However, the dense, highly domain-specific technical language currently used in these publications makes them inaccessible to the general public. This creates a significant barrier for potential donors and patients who wish to engage in the advancement of cancer research but feel they lack adequate knowledge. Additionally, it is important for those who have already contributed to research to be able to discover how their data has made an impact on our understanding of the disease.

Recent advances in large language models (LLMs) offer a promising solution to this problem. LLMs have demonstrated their abilities for text processing and summarization, and recent work has shown its applicability for texts in the medical domain^3,4^. When applied to simplify complex research articles at an appropriate level for the average person, these models can increase accessibility and promote informed participation in scientific research studies. Using LLMs to summarize medical-related texts is a goal shared by many and has proven successful in tasks like summarizing radiology reports^3^, patient symptom forms^5^ and hospital discharge information from electronic health records^6^. Interest in this area of engaging patients has also led to abstract-focused summarization of scientific literature^7^, but full-text manuscripts are rarely summarized, potentially due to challenges with LLM token limits, closed-access full text articles, and in validation with human-reviewed gold standard summaries which are time-intensive to generate. Existing work by Rinderknecht and colleagues^4^ summarized prostate cancer manuscripts and evaluated summaries by using automated metrics and by employing a single human expert rater to evaluate summary quality. We extended this approach to understand the accessibility and factuality of summaries by evaluating with a broader suite of metrics and a larger lay audience of varying scientific expertise.

In this study, we present a Python-based pipeline that uses PubMed to retrieve renal cancer research articles and prompts an LLM to generate lay language summaries from both the abstracts and full texts. We were also interested in *comparing* abstract-based summaries with full-text summaries using both automated evaluation methods and human-review by researchers, patient partners and lay audience members. Using quantitative metrics such as readability and grade scores, we show several metrics in agreement with human-review of the summaries, demonstrating that our approach is effective in summarizing medical texts.

### Approach

Using Metapub and Entrez^8,9^, open-access papers and their abstracts are retrieved, parsed, and subsequently summarized. As most of our work pertains to cancer research, we decided to focus our efforts in this domain. The scientific texts that are chosen are dictated by a keyword - for example “renal cancer” in our case - contained in a configuration file and modifiable by the user. The texts were then appended with a custom summarization prompt and provided to the chosen LLM. We selected the GPT-4-Turbo model based on previous work demonstrating the success of the latest GPT-based models in other medical-related summarization tasks^10,11^ and for its higher token limit, which more readily deals with the demanding task of summarizing full-text manuscripts. The “temperature” hyperparameter of the LLM was set to 0.5, reflecting recent work showing optimal instruction following and summarization while allowing for textual variability for our target lay audience^12^.

Our goal was to produce summaries that would be suitable for the general public. We aimed for a 6th to 8th grade reading level in order to ensure maximum readability, as advised by The National Institutes of Health (NIH) guidelines for health materials^13^ and Canada’s Research Ethics Board guidelines on consent forms^14^.

Multiple metrics were used to evaluate the performance of the model in generating lay summaries. These metrics include the Flesch Reading Ease score, the Flesch-Kincaid Reading Grade score, and a RAGAS Faithfulness score^15–17^. The Reading Ease and Grade scores were calculated by processing summaries using the Textstat Python library^18^ and the RAGAS score using the RAGAS Python library ^19^:

1. The Flesch Reading Ease score evaluates how comprehensible the text is by accounting for sentence length, word length, and number of words with more than 3 syllables. This value ranges from 0 to 100, with values closer to 100 indicating higher ease of reading.
2. The Flesch-Kincaid Reading Grade score is a measure of the United States education level for which the text is appropriate. Ranging from 0 to 18, we aimed for a value between 6 to 8, equivalent to a 6th to 8th grade literacy level.
3. The RAGAS Faithfulness metric^17^ measures if the generated summaries were factually accurate to the articles they were generated from. This method uses an LLM to query if the contextual information from the summaries are supported by the input articles’ text, with scores ranging from 0 to 1, with higher scores indicating contextual agreement.

Statistical testing to compare Abstract vs Full Text based summaries used the Wilcoxon Rank-Sum/Mann-Whitney U test. Initial experimentation revealed that prompts asking for a 6th or 8th grade level were too complex. To achieve a Flesch-Kincaid Reading Grade within grade 6-8, the LLM needed to be prompted to aim for a grade 2 audience (see Box 1.).

#### Box 1.

**Final prompt sent to the LLM**

Prompt:

“Summarize this article in 250 to 350 words for a 2nd grade lay audience using simple words. The tone should be casual. Avoid emotive or negative language such as battle or fight. Be realistic in your language around impact and expected outcomes.”

In addition to automatic evaluations, we conducted a human review with 18 volunteers reviewing a subset of the 106 generated summaries, totalling 40 human reviewed summaries (Fig. 1). These 40 summaries were selected at random, and of these, 20 summaries were based on abstracts and 20 on full texts, with abstract-based summaries derived from different manuscripts to the full-text-based summaries. We used this procedure to accommodate the size of our human review group, to ensure each summary had a unique group of multiple reviewers, and to widen the range of manuscripts for which derived summaries could be reviewed. Each summary was independently reviewed by a unique group of 2 to 4 reviewers. Five reviewers were patient partners who were compensated for 3 hours of their time. The remaining volunteers were employees of the Ontario Institute for Cancer Research with varying degrees of biology background knowledge. Reviewers were asked to evaluate 5-7 summaries each via an evaluation form provided to them (see Box 2.). Three qualities were evaluated: readability, clarity, and accuracy. Each quality was defined at the beginning of the form, alongside a guide on how to score grade level. We also had each reviewer assess how equipped they felt to evaluate these texts, on a scale of 1 to 5. This metric allowed us to compare reviewers’ analyses of the summaries alongside their self-assessed experience level.

**Figure 1.**
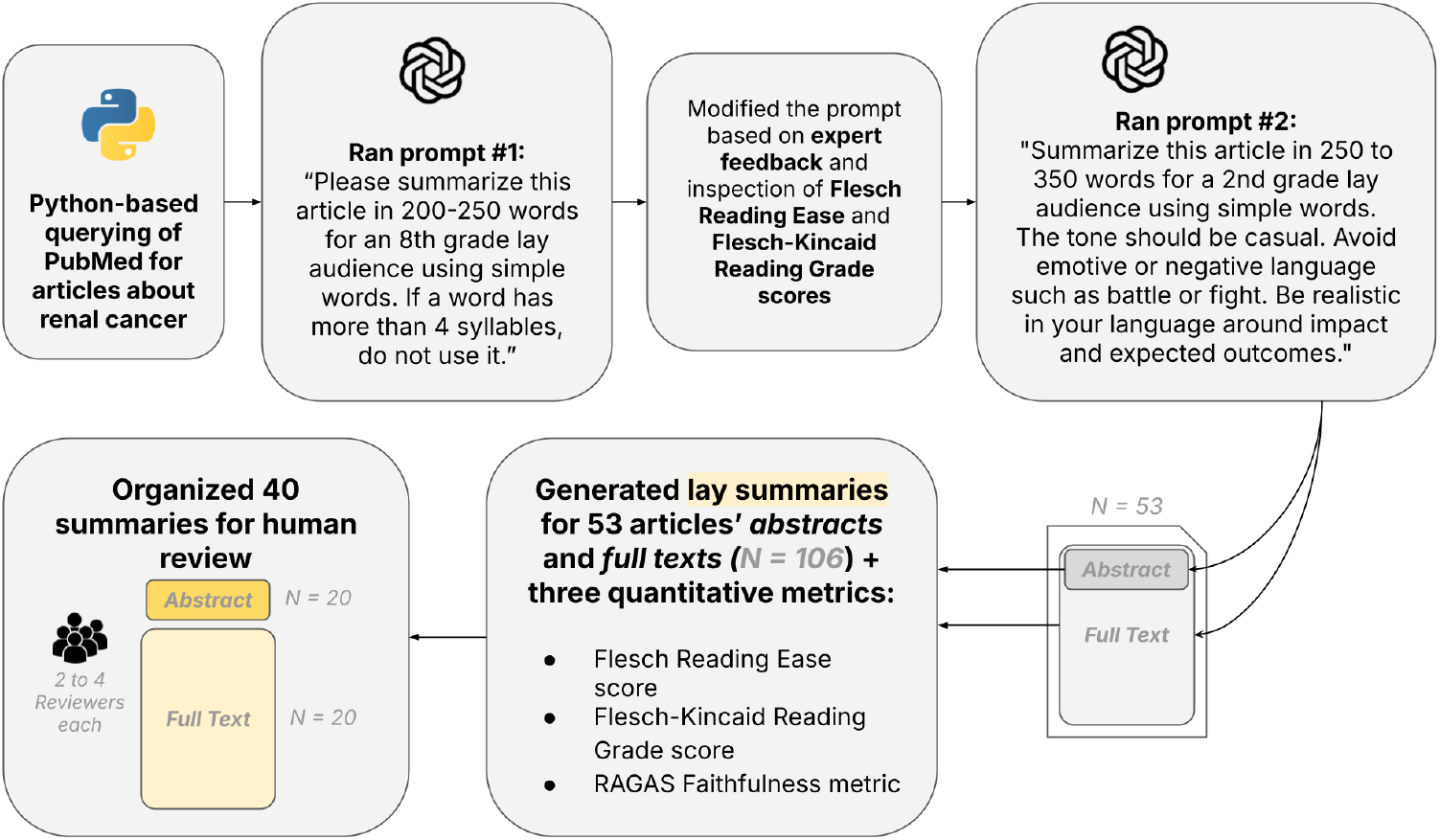
Study workflow detailing the process of summary generation, prompt modifications and evaluation for LLM-based lay summary generation. Evaluations were performed using both automated quantitative metrics, and human review.

#### Box 2.

**Example of the questions included in the form sent to human validation volunteers**

**SUMMARY #1**

Please provide a 1-2 sentence overview of Summary #1. (in your own words, describe what the summary intended to say)

How would you rate the clarity of summary #1? (1-10)

What reading grade level would you say summary #1 is appropriate for? (Grade 1-10)

How well did Summary #1 capture the key points of its corresponding article (either abstract or full-text)? (1-10)

Did lay summary #1 avoid jargon and technical terms appropriately? (Yes/No)

**CLOSING QUESTIONS**

On average, how effective were the summaries in simplifying academic renal cancer papers for a lay audience? (1-10)

Were there any factually false or misleading statements in the summaries you read? If no, write “No” and if yes, please specify them below.

Did you feel well-equipped to complete the task of reviewing the research papers and evaluating their lay summaries? (e.g. in terms of familiarity reading scientific literature) (1-5)

Patient partners were also asked to complete a post-evaluation survey, to understand their thoughts on the impact AI-generated lay summaries may have on participation in and understanding of research studies. Questions pertained to whether having access to AI-generated summaries of research findings would make them more likely to participate in future studies, as well as advantages and skepticisms they might have with respect to the effectiveness of AI-generated summaries.

## Results and discussion

We ran the pipeline, generating a total of 106 summaries – 53 abstract and 53 full-text. Querying PubMed for open access papers using the provided keyword “renal cancer”, GPT-4-Turbo summarized the results within the parameters dictated by the prompt. The automated summarization pipeline completed in minutes, a task that could require hours of manual effort by human experts.

The average Abstract Flesch Kincaid Reading Grade Score was 7.13 (*SD*=1.45) and the average Full Text Flesch Kincaid Reading Grade Score was 7.39 (*SD*=1.37) indicating summaries at around 7th to 8th grade reading level with no significant differences between Abstracts and Full Texts (*p*=0.39).

**Table 1.** Summary statistics from the results of 53 abstract summaries and 53 full text summaries. These automated metrics highlight that both the abstracts and full texts produce summaries that are readable according to thresholds set forth by the NIH and Health Canada, and faithful to their original texts.

|  | **Abstract** |  |  | **Full Text** |  |  |
| --- | --- | --- | --- | --- | --- | --- |
|  | N=53 |  |  | N=53 |  |  |
|  | Flesch Reading Ease | Flesch-Kincaid Grade Score | Faithfulness Score | Flesch Reading Ease | Flesch-Kincaid Grade Score | Faithfulness Score |
|  | *0-100, Higher is better* | *Target: 6-8* | *0-1, Higher is better* | *0-100, Higher is better* | *Target: 6-8* | *0-1, Higher is better* |
| **Mean** | 73.21 | 7.13 | 0.84 | 72.07 | 7.39 | 0.88 |
| **Median** | 72.62 | 7.19 | 0.89 | 71.52 | 7.32 | 0.92 |
| **Standard Deviation** | 5.98 | 1.45 | 0.17 | 5.43 | 1.37 | 0.13 |
| **Minimum** | 58.71 | 4.07 | 0.13 | 59.71 | 4.31 | 0.53 |
| **Maximum** | 86.08 | 11.44 | 1.00 | 84.23 | 10.17 | 1.00 |

The Flesch Reading Grade Level score places heavy emphasis on length of sentences and number of syllables per word. Given the medical and academic context of these lay summaries, it is difficult to avoid multi-syllabic words, such as “carcinoma” or “chemoradiotherapy”. Notably, LLM-generated summaries often provided brief explanations for abbreviations and biomedical jargon such as “mRNA” or “autophagy”, which may serve to improve understanding of these multi-syllabic words. While the Flesch Reading Grade Level provides a standard score to compare summaries from previous works, unlike its rigidity and focus on word complexity, LLMs have the capacity to appraise the semantics of longer text sequences and acknowledge that the summaries are effectively simplifying technical contexts. We fed the summaries back through GPT-4 and asked “What grade level is this text?”. The LLM could recognize that the summaries’ topics were advanced while still evaluating the simplicity of language and analogies. This qualitative analysis of the summaries by the LLM proved to be more aligned with the grade level assessments by human review, while also yielding results that were, as expected, quite different from the Flesch-Kincaid Reading Grade Level (Fig. 3).

Previous work has shown the impact of prompt engineering on improving lay summaries readability and grade level suitability^4^. To study the impact of prompt complexity and instruction, a simpler alternative prompt was tested: *“Please summarize this article in 200-250 words for an 8th grade lay audience using simple words. If a word has more than 4 syllables, do not use it*.*”* With this prompt, the average Flesch Grade Level scores were 9.58 (abstract) and 11.78 (full text) and the Reading Ease scores were 58.61 (abstract) and 43.78 (full text). As expected, our aim to achieve appropriate reading grade level scores around 8th grade as opposed to the 12th-14th grade levels achieved by Shyr et al. (2024) yielded significantly higher reading ease scores. Though, expert feedback of summaries generated with this prompt noted that the summaries should be longer, the language and tone should be casual and conversational, we should avoid emotive language, and the grade level should be lower. These comments inspired the final prompt, which incorporates details to support this feedback. A prompt providing more detailed instructions reduced the median grade level from 14.8 to 12.8, while increasing reading ease scores from 29.1 to 40.9, a trend consistent with our metrics when prompt modifications were implemented.

Despite the prompt requesting lay summaries appropriate for a 2nd grade audience, the resulting Flesch-Kinkaid grade score metrics were on average much higher (7.13 for abstract and 7.39 full text). This finding reinforces the notion that there is a disparity between the level specified in the prompt and the yielded summaries’ grade level calculated using the Flesch-Kincaid Grade Level score (Supplementary Fig. 1). This discovery emphasized the importance of qualitative analyses of these summaries to discern their simplicity from a human perspective.

Feedback provided by human reviewers indicated that the tone of some summaries was “condescending,” “juvenile,” or “overly simplified.” The average clarity and accuracy scores were 7.41/10 and 7.23/10 respectively. When asked if jargon was minimized appropriately, 13% of reviewers said “No,” with some reviewers expressing that jargon was minimized *too* much. This raises questions about whether the quantitative metrics’ can appropriately gauge tone, language, and vocabulary, and in turn their ability to determine if a text is actually suitable for its stated audience. While the prompt requested the summaries be written at a 2nd grade level, the average full text Flesch-Kincaid Grade level was 7.39 and the average grade level provided by human reviewers was 5.75. While there exists a disparity between these two results, they are nonetheless both significantly above the 2nd grade level specified in the prompt, and closer to NIH/Health Canada recommendations of a 6th-8th grade level. Future work may address the unsuitable tone of the summaries with a prompt that emphasizes summaries for “an adult audience with 2nd grade level reading comprehension”, as opposed to our current prompt which may imply a lay audience of 2nd grade age, and specifies using “simple words”.

When comparing scores based on human reviewers’ self assessment of expertise (Fig. 4), median reviews of accuracy and clarity in the highest expertise level gave similar scores to that in the lower expertise levels (8 for expertise level 2 and 5). Median grade level ratings were slightly lower with greater expertise (grade 5 from expertise level 4 and 5 vs grade 7 in expertise level 3). While there are limitations to objectivity for these self reported measures of expertise, they do provide confirmation that the quality of summaries is fairly consistent across a wide audience of readers. Additionally, human review of accuracy only showed a slight, non-significant improvement when generating from full text versus abstracts (Supplementary Fig. 2), aligning with the automated RAGAS metrics results (Fig. 2C). This may indicate that abstract-based LLM summarization may be sufficient, providing cost-savings particularly in low resource settings where a greater number of LLM tokens would be required for full-text summarization.

**Figure 2.**
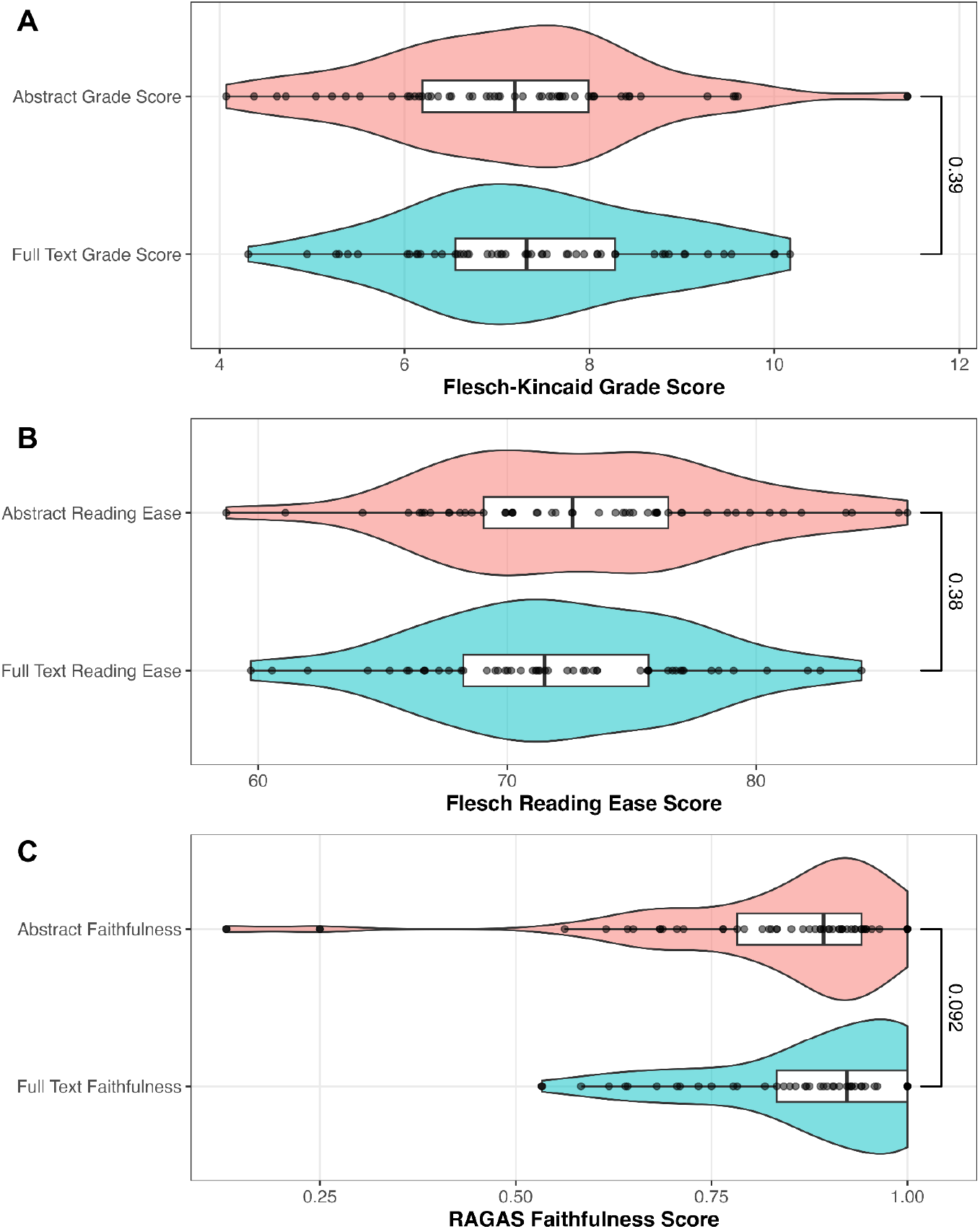
Comparison between the performance of the abstract and full-text summaries given 3 quantitative metrics: Flesch-Kincaid Grade Score, Flesch Reading Ease, and RAGAS Faithfulness. No significant differences were found between summaries generated from abstracts versus full-texts in the Flesch-Kincaid Grade Score, Flesch Reading Ease or RAGAS Faithfulness (*p=0*.*39, 0*.*38, 0*.*092* respectively).

**Figure 3.**
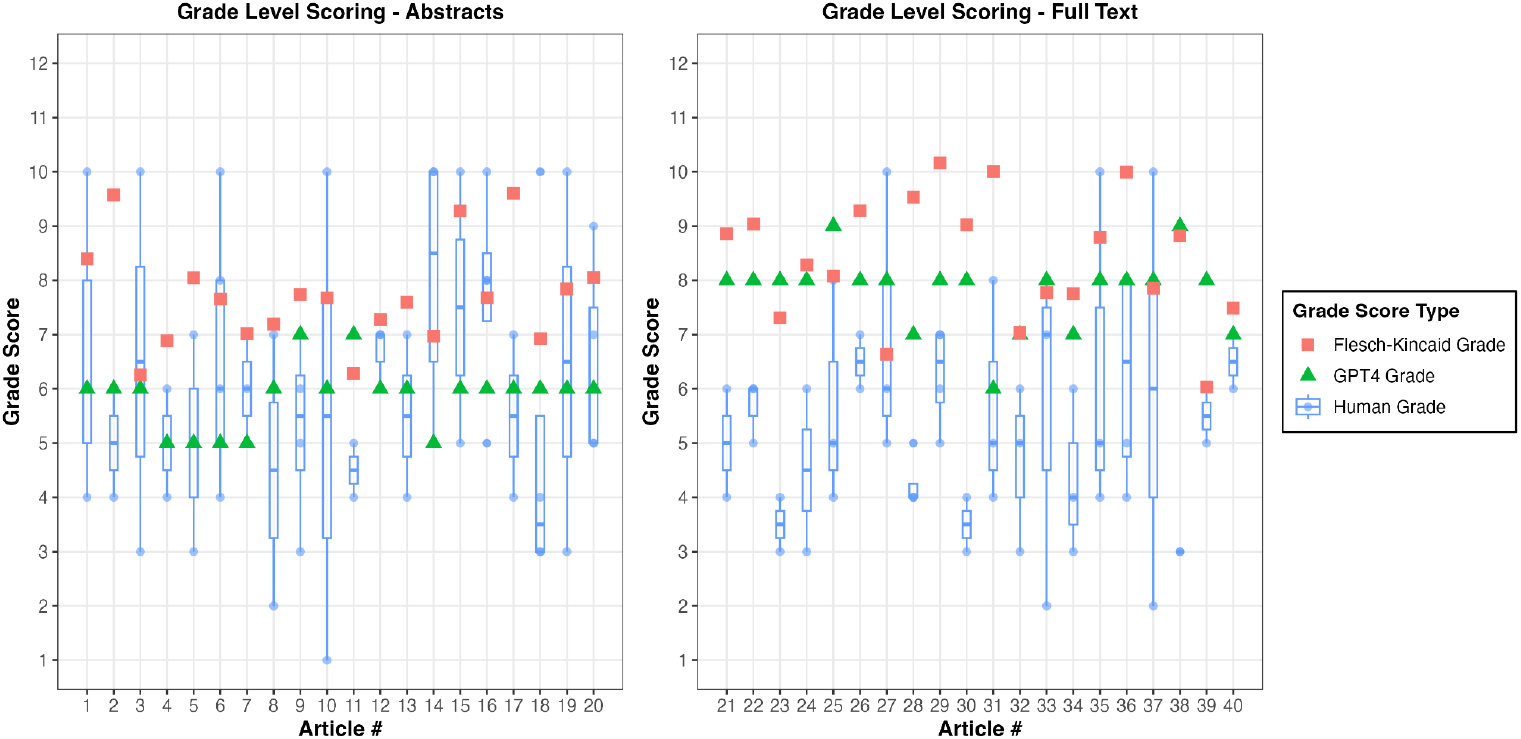
Average Grade Level scores generated by the Flesch-Kincaid Reading Grade Score, GPT-4 self-assessment, and human reviewers for abstract (left) and full-text (right) lay summaries for the subset of 40 articles’ summaries that were human reviewed. Flesch-Kincaid Grade Levels exceed the median human review grade level for 37 (92.5%) of the articles, and GPT-4 grade level appraisals for 33 articles (82.5%).

**Figure 4.**
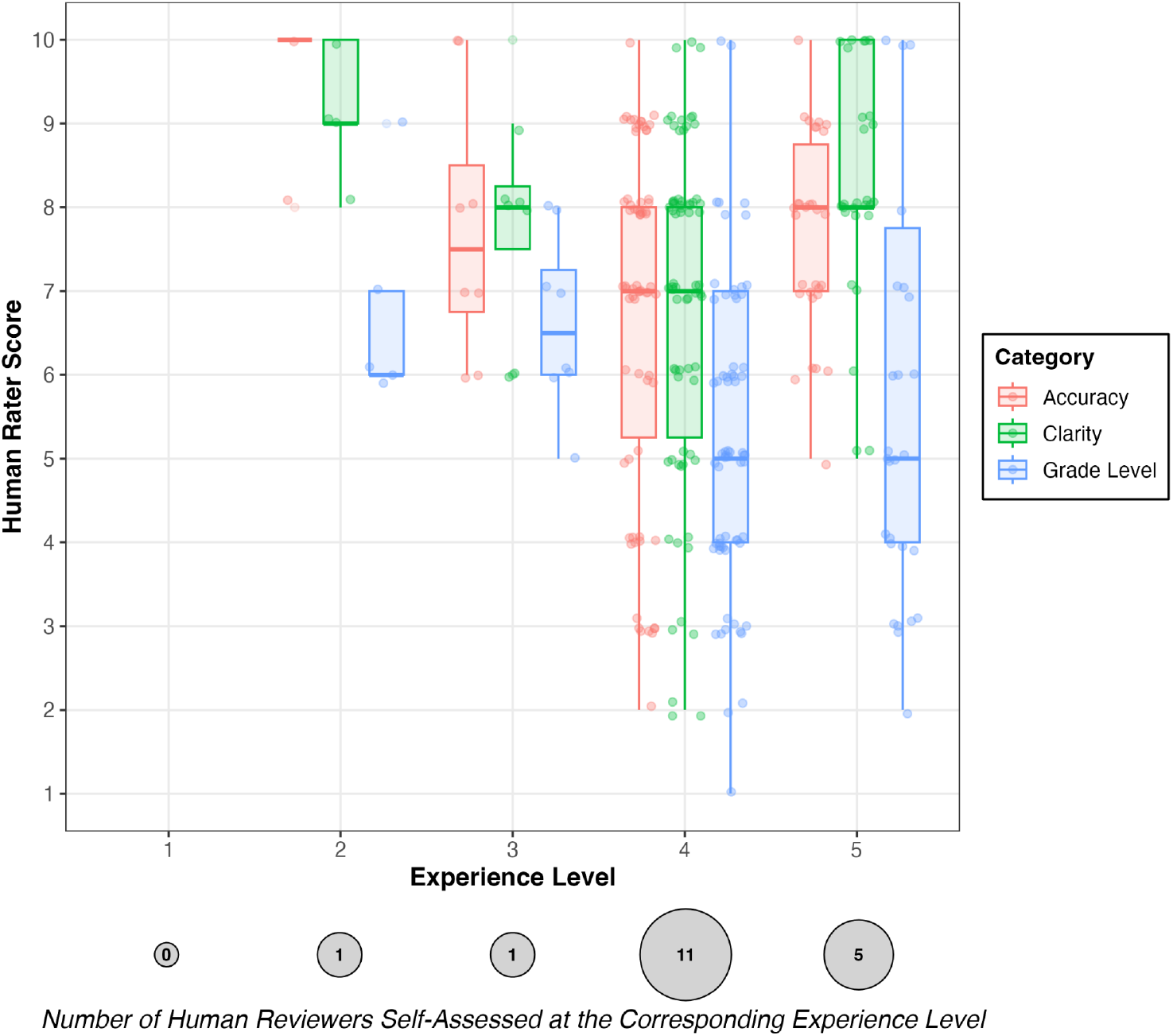
Human rating counts of accuracy, clarity and grade level scores grouped on self-reported experience level (1 through 5) of human reviewers.

Although the grade level classification as judged by human reviewers met the recommended target, we find that specifying grade level in the prompt is not effective in generating appropriate lay summaries. In addition to adopting the vocabulary or sentence structure of an elementary audience, the results demonstrate that the summaries were too simple at times, even hesitating to use terms like “genes” or “cancer.” A future approach may consist of listing out the qualities we had desired, such as minimal jargon and short sentences, but without the 7-8 year old age group that is associated with 2nd grade.

In the patient partner post-evaluation survey, all partners responded that they would find such an AI-generated lay summary to be useful for their understanding and for guiding their decision to participate in a research study (Fig. 5); all patients partners mentioned the summary would make them more willing to enroll. One patient mentioned removing “overwhelming” and “complex” language was helpful for making them feel informed, and overall “patient impact” and “practical implications” of studies was emphasized to be important. It was also evident that while our approach could reduce a barrier patients face with complex scientific writing, skepticism with the reliability of AI remains. Credibility of sources and secondary human review of the medical texts were factors that several patients noted would help reinforce their trust in the summaries.

**Figure 5.**
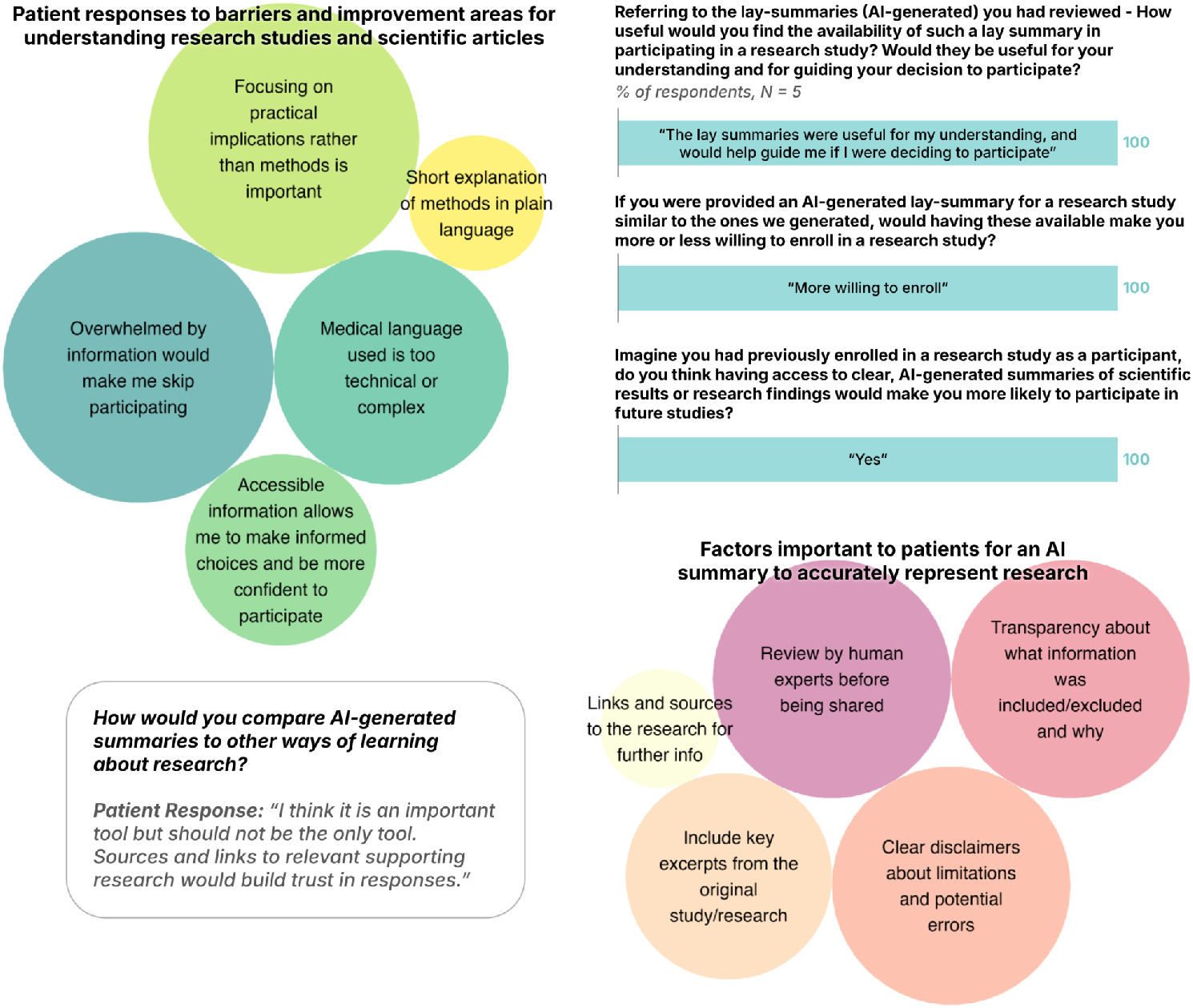
Patient partner post-evaluation survey results. Patient partners provided perspectives on their views as to how automatically generated lay summaries might increase participation and engagement in research studies through improved understanding. Two core themes emerged: 1) Ensuring summaries’ trustworthiness and transparency; and 2) maintaining patient-centric language. Size of bubbles are proportional to the number of times themes in the embedded text excerpts were mentioned in the survey responses.

### Limitations

This study has several important limitations to consider. First, even with LLMs set to a minimal “temperature”, the stochastic nature of LLMs means that generated summaries can vary subtly between iterations, potentially affecting the language and content included in summaries. This inherent variability suggests that further validation with a larger set of medical or scientific texts would strengthen confidence in the trends we uncovered, and provide additional opportunity to examine the ability of summary generation to be personalized and defined to an LLM according to user-preferred reading level, tone and language.

As our analysis focused on participants’ feedback, it was constrained by a relatively small sample size of only 40 articles’ summaries for which we obtained both automatic metrics and human evaluations. While this sample size did provide meaningful insight into the relative performance and perceived qualities between abstract vs full-text-based summaries, a large-scale head-to-head comparison within individual articles would extend our findings and complement our validation approach. We also focused human review metrics (such as accuracy) of abstracts versus full-text in 2 distinct subsets of 40 articles rather than a smaller head-to-head approach to broaden the coverage of manuscripts’ subject matter given the limited size of our human review cohort. Using this approach on a larger human review cohort and article group would further validate these patterns of summary perception and quality across a more diverse set of scientific material.

Additionally, any human evaluation process inherently involves subjective judgment, for instance in our study the perceived accuracy of summaries. Our reviewers likely possessed higher-than-average scientific expertise and comprehension levels, which enabled a more rigorous evaluation of how the scientific content was represented. However, this level of familiarity may differ from the preferences of general lay audiences who are the intended readers of these summaries, and our lay summaries alone will not be able to assist lay audiences in judging the scientific merit of a publication. Understanding how an even more diverse audience of varying expertise levels responds to AI-generated summaries represents a valuable next step for future research in improving science communication.

## Conclusion

Through this work, we aim to bridge the gap between jargon-heavy scientific literature and lay summaries suitable for the average reader. By exploring the use of LLMs to generate lay summaries of renal cancer papers, our tool drastically streamlines summarization and operates at a fraction of the time as manual human summarization alone. Our approach demonstrates that GPT-4-Turbo can produce summaries that human reviewers found to be readable and faithful to their original texts. These findings suggest that AI-generated summaries have promising potential when it comes to bridging the gap between scientific communities and the general public. Future work should investigate custom training and fine-tuning strategies to improve performance in a variety of different medical contexts.

Our lay summary pipeline will ultimately benefit patients by making complex biomedical research more accessible and understandable. By translating full-text scientific articles into clear, accurate, and easy-to-read summaries, patients and their families can better engage with the latest medical findings and make more informed decisions about their health. This tool will serve as one component to improving information accessibility, and we hope it can be used in conjunction with additional patient engagement strategies to expand the way the research community fosters connection with the public.

## Acknowledgements

We thank Aaron Yu, Andres Melani de la Hoz, Barbara Dresner, Edmund Su, Harjeet Kaur, Katarina Maksimovic, Linda Xiang, Lindsey Wolfram, Lucy Ioannoni, Mitchell Shiell, Paige Johnson, Sarah Barker, Sarah Ridd and other members of our human reviewer group for their time with the lay summary evaluation. Special thanks to Robin Haw for his feedback on initial summaries, and Justin Noble for his help with recruiting patient partners.

## Code and Data Availability

Code for the pipeline and figures can be found on GitHub. (https://github.com/courtotlab/Lay-Summary-Project). Summaries and data to reproduce the figures can be found on Figshare (doi:10.6084/m9.figshare.30827678). Generative Artificial Intelligence (AI) models including GPT-4 within Github CoPilot were used to assist the development of code for the data processing pipelines. After using the tool, authors reviewed and modified the code, and take full responsibility for the content of the publication.

## Supplementary Figures

**Supplementary Figure 1.**
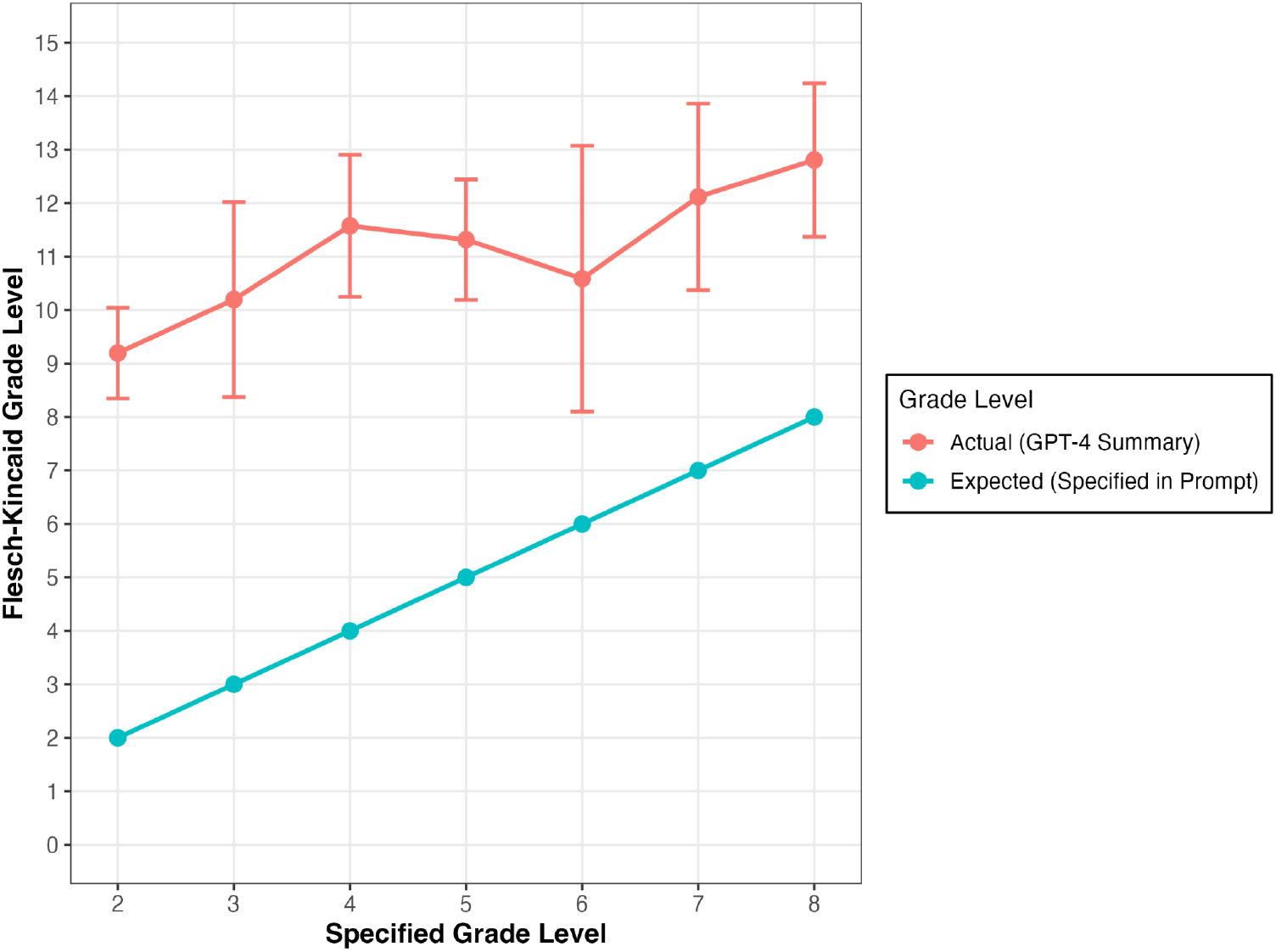
Relationship between the grade level specified in the prompt (Expected, refer to Box 1 for prompt) and mean Flesch-Kincaid grade level scores of generated summaries (Actual). Actual grade level means and standard deviation is plotted alongside the expected grade levels. Actual grade levels exceed the expected levels requested, at all grade levels.

**Supplementary Figure 2.**
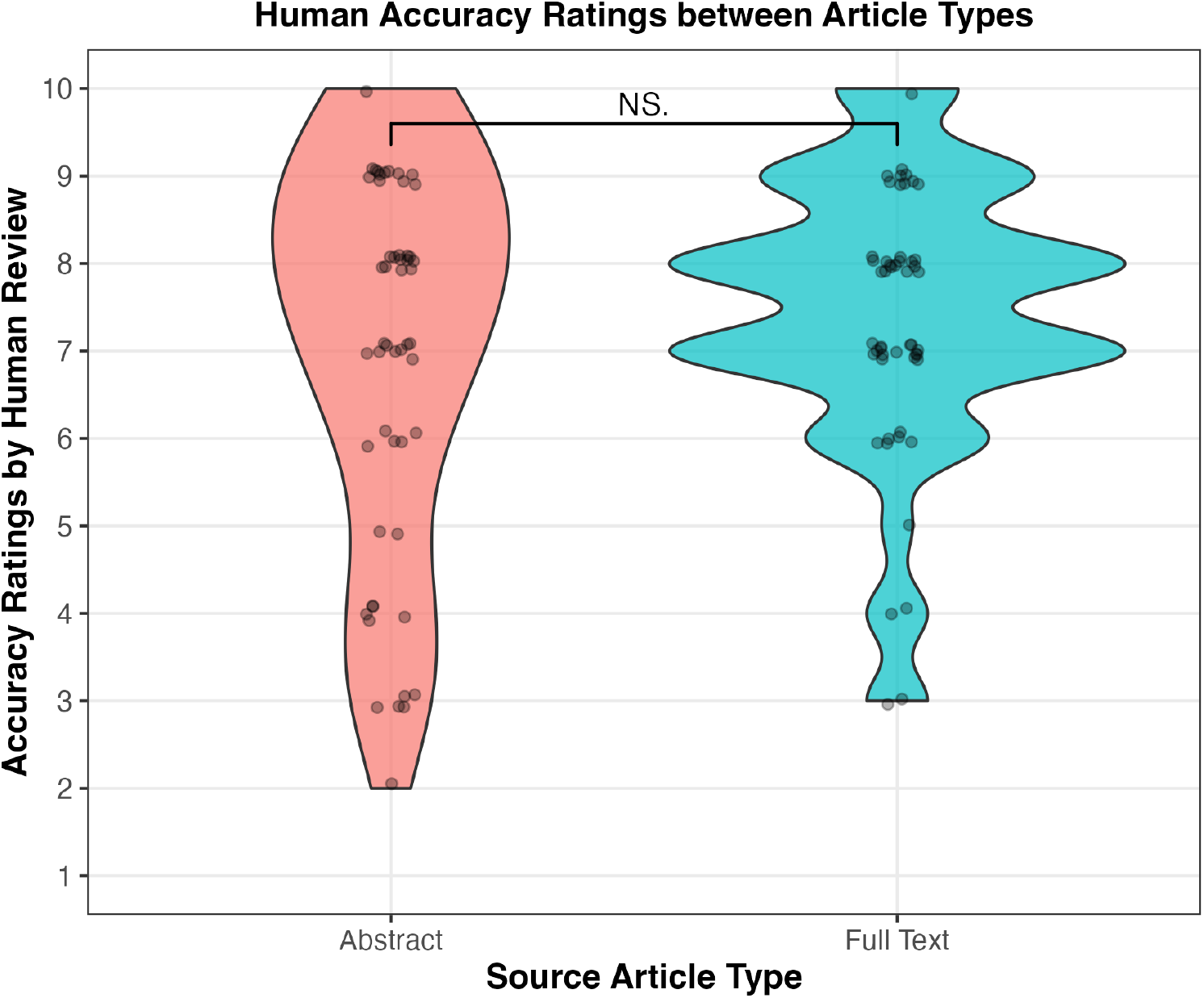
Accuracy ratings (0-10) from human review of summaries generated from abstracts compared to those generated from full text articles.

